# Suppression of *cdc13-2*-associated senescence by *pif1-m2* requires Ku-mediated telomerase recruitment

**DOI:** 10.1101/2021.03.31.437837

**Authors:** Enikő Fekete-Szücs, Fernando R. Rosas Bringas, Sonia Stinus, Michael Chang

## Abstract

In *Saccharomyces cerevisiae*, recruitment of telomerase to telomeres requires an interaction between Cdc13, which binds single-stranded telomeric DNA, and the Est1 subunit of telomerase. A second pathway involving an interaction between the yKu complex and telomerase RNA (TLC1) contributes to telomerase recruitment, but cannot sufficiently recruit telomerase on its own to prevent replicative senescence when the primary Cdc13-Est1 pathway is abolished—for example, in the *cdc13-2* mutant. In this study, we find that mutation of *PIF1*, which encodes a helicase that inhibits telomerase, suppresses the replicative senescence of *cdc13-2* increasing reliance on the yKu-TLC1 pathway for telomerase recruitment. Our findings reveal new insight into telomerase-mediated telomere maintenance.

## Introduction

Telomeres are composed of G/C-rich repetitive sequences at the termini of eukaryotic chromosomes and play a pivotal role in genome maintenance by “capping” chromosome ends, preventing them from unwanted nucleolytic degradation, homologous recombination, and fusion with neighboring chromosomes (Jain & Cooper, 2010). In addition, to overcome progressive telomere shortening due to the end replication problem, telomeres are elongated by a specialized reverse transcriptase called telomerase. In the budding yeast *Saccharomyces cerevisiae*, telomerase is minimally composed of the protein subunit Est2 and the RNA subunit TLC1 (Lingner *et al*, 1997; Singer & Gottschling, 1994). However, additional accessory proteins, Est1 and Est3, are required for telomerase activity *in vivo*, and are thought to be involved in the recruitment and/or activation of telomerase (Wellinger & Zakian, 2012). Eliminating any of the Est proteins or TLC1 results in an “ever shorter telomeres” (*est*) phenotype characterized by progressive telomere shortening that ultimately leads to replicative senescence (Lundblad & Szostak, 1989; Lendvay *et al*, 1996; Singer & Gottschling, 1994).

Maintaining telomere length homeostasis through the regulation of telomerase is essential for genome stability. Several lines of evidence suggest that the recruitment of telomerase to telomeres involves a direct interaction between the Est1 subunit of telomerase and Cdc13, a protein that binds single-strand telomeric DNA with high affinity (Lin & Zakian, 1996; Nugent *et al*, 1996). Expression of a Cdc13-Est2 fusion protein can support telomere maintenance in an *est1Δ* null mutant, suggesting that the main function of Est1 is to bring telomerase to telomeres (Evans & Lundblad, 1999). Cdc13 is essential for telomere capping, so a null mutation is lethal; however, an extensively studied point mutant, *cdc13-2*, is not capping defective, but displays an *est* phenotype (Nugent *et al*., 1996). The amino acid mutated in *cdc13-2*, E252, lies within the recruitment domain (RD), which is able to recruit telomerase to telomeres when fused to the DNA binding domain of Cdc13 (Pennock *et al*, 2001). The mutation (E252K) results in a charge swap and can be suppressed by *est1-60*, which encodes a mutant Est1 with a reciprocal charge swap (K444E), suggesting a direct physical interaction between the two proteins (Pennock *et al*., 2001). Consistent with this idea, purified full-length Cdc13 and Est1 interact *in vitro* (Wu & Zakian, 2011), and structural analysis revealed two conserved motifs within the Cdc13 RD, called Cdc13_EBM-N_ and Cdc13_EBM-C_ (referring to N- and C-terminal Est1-binding motifs, respectively), responsible for this interaction (Chen *et al*, 2018). The Cdc13 E252K mutation resides within the latter motif. Surprisingly, mutations in the Cdc13_EBM-C_ motif, including E252K, do not abolish the interaction between Cdc13 and Est1 *in vitro* despite causing a dramatic reduction in Est1 telomere association *in vivo* (Wu & Zakian, 2011; Chen *et al*., 2018; Chan *et al*, 2008). Thus, the mechanism by which the Cdc13_EBM-C_ motif promotes telomerase-mediated telomere extension is still unclear.

In contrast, mutations in Cdc13_EBM-N_ abolish the Cdc13-Est1 interaction *in vitro*, yet only result in a modest reduction in Est1 telomere association and short, but stable, telomere length *in vivo* (Chen *et al*., 2018). This telomerase recruitment pathway works in parallel with a second pathway involving Sir4, the yKu complex, and TLC1. Doublestrand telomeric DNA is bound by Rap1 (Buchman *et al*, 1988; Conrad *et al*, 1990), which interacts with Sir4 (Moretti *et al*, 1994). Sir4, in turn, interacts with the Yku80 subunit of the yKu complex (Roy *et al*, 2004), which binds to the tip of a 48-nt hairpin in TLC1 (Peterson *et al*, 2001; Stellwagen *et al*, 2003; Chen *et al*., 2018). Mutations that abolish the yKu-TLC1 interaction (e.g. *tlc1Δ48* and *yku80-135i*) have slightly short but stable telomeres (Peterson *et al*., 2001; Stellwagen *et al*., 2003), much like Cdc13_EBM-N_ mutations. Disrupting both the yKu-TLC1 interaction and Cdc13_EBM-N_-Est1 interaction results in an *est* phenotype (Chen *et al*., 2018).

Pif1, a 5’-3’ helicase that is evolutionary conserved from bacteria to humans, directly inhibits telomerase activity at telomeres and DNA double-strand breaks (Schulz & Zakian, 1994). Pif1 has both mitochondrial and nuclear isoforms; by altering the first (*pif1-m1*) and the second (*pif1-m2*) translational start sites, the functions can be separated (Schulz & Zakian, 1994). The *pif1-m2* mutant abolishes nuclear Pif1 and, similar to *pif1Δ*, has elongated telomeres (Schulz & Zakian, 1994). *In vitro*, purified Pif1 reduces telomerase processivity and displaces telomerase from telomeric oligonucleotides (Boulé *et al*, 2005). *In vivo*, deletion of *PIF1* increases telomere association of Est1, while overexpression of *PIF1* reduces telomere association of Est1 and Est2 (Boulé *et al*., 2005).

We previously showed that a double-strand break adjacent to at least 34 bp of telomeric sequence is efficiently extended by telomerase, resulting in the addition of a *de novo* telomere, but this does not occur in Cdc13_EBM-C_ mutants, such as *cdc13-2* (Strecker *et al*, 2017). Surprisingly, we found that the lack of telomere addition in *cdc13-2* cells can be suppressed by the *pif1-m2* mutation (Strecker *et al*., 2017). In this study, we find that *pif1-m2* suppresses the replicative senescence caused by the *cdc13-2* mutation in a manner dependent on the yKu-TLC1 telomerase recruitment pathway. In addition, *pif1-m2* suppresses the replicative senescence caused by disrupting both the yKu-TLC1 and Cdc13_EBM-N_-Est1 interactions. These observations provide new insight into the complexity of telomerase-mediated telomere maintenance.

## Results and Discussion

### Mutation of *PIF1* suppresses the replicative senescence caused by the *cdc13-2* mutation

To investigate how telomere addition is possible in a *cdc13-2 pif1-m2* genetic background, we serially passaged cells to determine whether they would senesce. For these experiments, *cdc13Δ* or *cdc13Δ pif1-m2* cells, kept alive by the presence of a high-copy plasmid expressing wild-type *CDC13* and the *URA3* gene, were transformed with an additional plasmid containing either *CDC13* or *cdc13-2*. These cells also carried a deletion of *RAD52* to prevent homologous recombination-mediated telomere maintenance (Claussin & Chang, 2015). We then counterselected the first plasmid by growing cells on media containing 5-fluoroorotic acid (5-FOA), which is toxic to cells expressing *URA3*. 5-FOA-resistant colonies were subsequently serially passaged on agar plates (Fig 1A). Senescence was apparent for *cdc13-2 PIF1* cells already after the first passage, whereas *CDC13* and *cdc13-2 pif1-m2* strains did not show any sign of senescence even after the fourth passage. We analyzed the telomere length of these strains and found that, consistent with previous studies, *pif1-m2* has increased telomere length compared to wild type (Schulz & Zakian, 1994) while the telomeres are very short in the *cdc13-2* mutant (Lendvay *et al*., 1996; Nugent *et al*., 1996). Interestingly, *cdc13-2 pif1-m2* telomeres are approximately wild-type in length, albeit more heterogeneous, and stable throughout the course of the experiment (Fig 1B). Our findings indicate that telomerase-mediated telomere extension can occur in *cdc13-2 pif1-m2* cells, allowing cells to maintain telomere length homeostasis and avoid replicative senescence.

**Figure 1.**
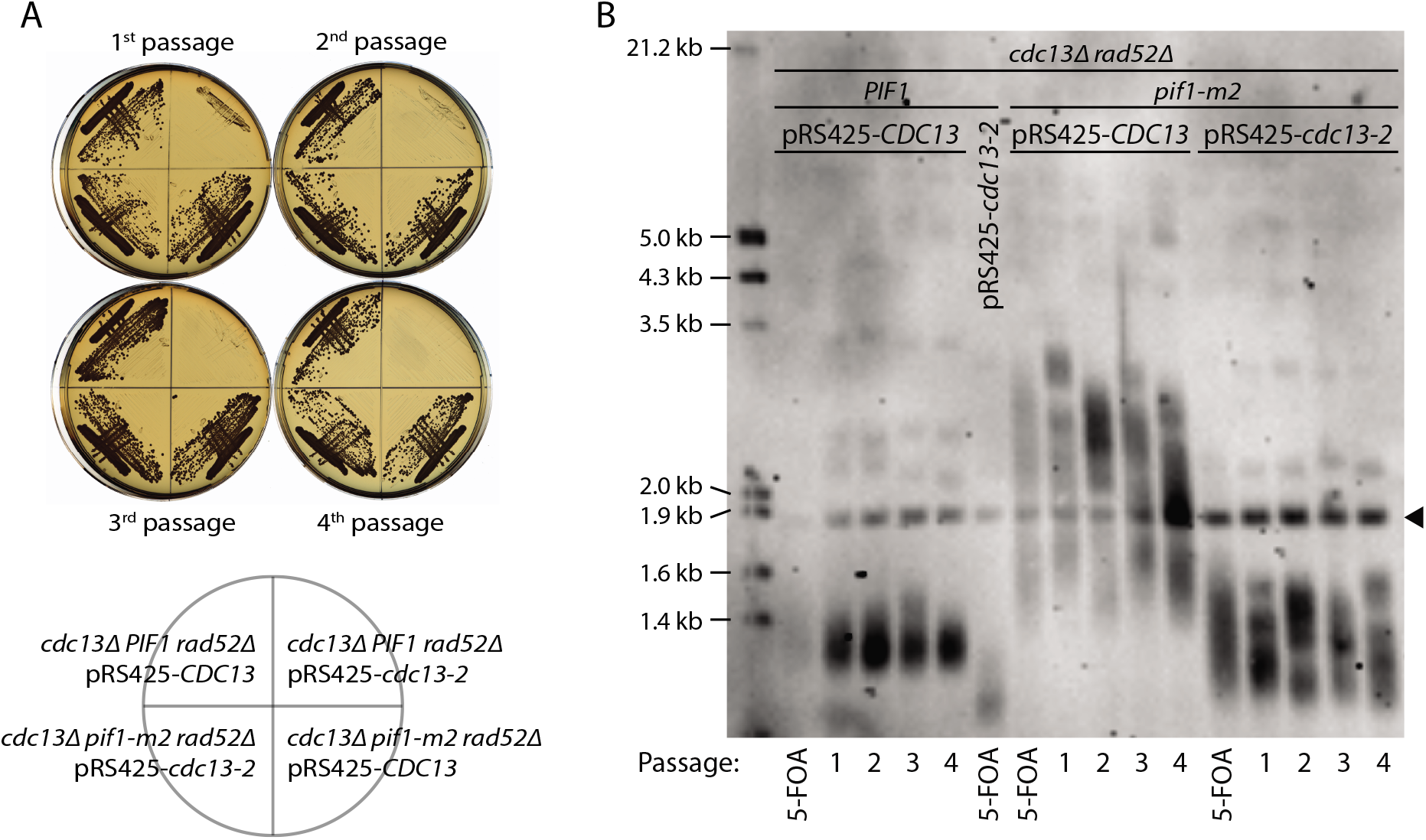
*cdc13-2 pif1-m2* cells do not senesce and have telomeres that are stable in length. A Strains of the indicated genotypes were passaged four times on YPD plates after counterselection on 5-FOA to remove plasmid YEp24-*CDC13*. Each passage corresponds to ~25 generations of growth. B Telomere Southern blot analysis of strains from A. The black arrowhead indicates a 1.8 kb DNA fragment generated from the BsmAI-digestion of plasmid pYt103.

### The yKu-TLC1 telomerase recruitment pathway is necessary to maintain telomere length in *cdc13-2 pif1-m2* cells

We hypothesized that the yKu-TLC1 pathway may become essential for telomere length homeostasis in *cdc13-2 pif1-m2* strains. To test this possibility, haploid meiotic progeny derived from the sporulation of *CDC13/cdc13-2 PIF1/pif1-m2 YKU80/yku80-135i* and *CDC13/cdc13-2 PIF1/pif1-m2 TLC1/tlc1Δ48* heterozygous diploids were serially propagated in liquid culture for several days (Fig 2A and 2B). The *yku80-135i* and *tlc1Δ48* alleles disrupt the interaction between the yKu complex and TLC1 (Peterson *et al*., 2001; Stellwagen *et al*., 2003). As expected, *cdc13-2* cultures grew slower as the experiment progressed and cells senesced, but growth was eventually restored upon the emergence of survivors that utilize recombination-mediated mechanisms to maintain telomeres (Lendvay *et al*., 1996). In contrast, the *cdc13-2 pif1-m2* strains did not senesce, confirming our previous observations (Fig 1). The *cdc13-2 pif1-m2 yku80-135i* and *cdc13-2 pif1-m2 tlc1Δ48* triple mutants showed a pattern of senescence and survivor formation, indicating that the yKu-TLC1 telomerase recruitment pathway is required for telomere length homeostasis in *cdc13-2 pif1-m2* cells. The *yku80-135i* and *tlc1Δ48Δ* alleles caused *cdc13-2* and *cdc13-2 pif1-m2* strains to senesce faster, but the reason for this is currently unclear.

**Figure 2.**
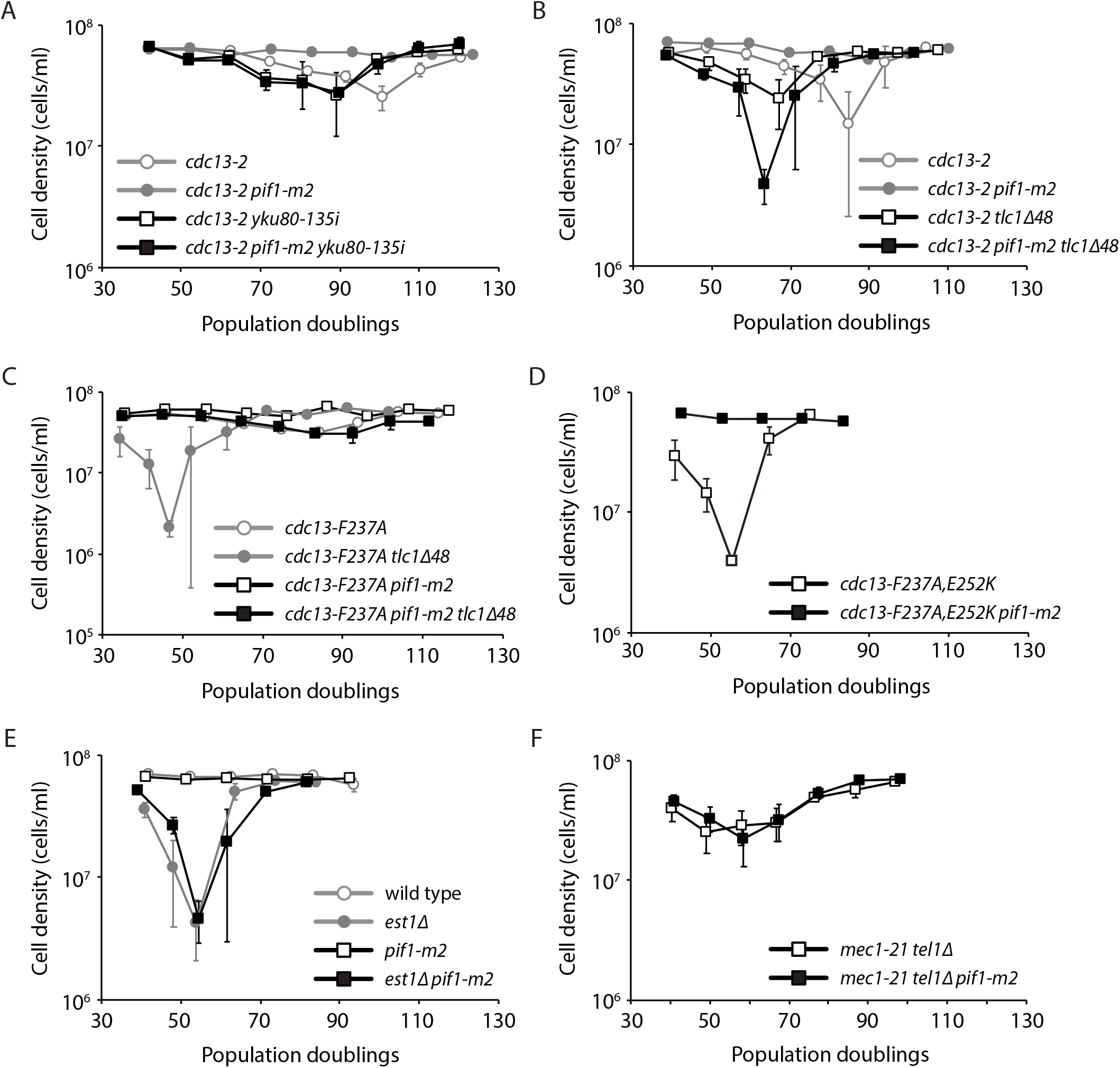
Telomeres are maintained by the yKu-TLC1 pathway in *cdc13-2 pif1-m2* cells. Senescence was monitored in liquid culture by serial passaging of haploid meiotic progeny derived from the sporulation of VSY20 (A), VSY7 (B), EFSY73 (C), FRY867 (D), CAY2 (E), and MCY815 (F). Average cell density ±SEM of 3-4 independent isolates per genotype (except n=2 for *cdc13-F237A, E252K*) is plotted.

Combining mutations that disrupt the Cdc13_EBM-N_-Est1 interaction (e.g. *cdc13-F237A*) and the yKu-TLC1 interaction leads to replicative senescence (Chen *et al*., 2018). We tested whether the *pif1-m2* mutation could suppress this replicative senescence and found that it can: *cdc13-F237A tlc1Δ48* strains senesce while *cdc13-F237A pif1-m2 tlc1Δ48 strains* do not (Fig 2C). Similarly, *pif1-m2* can suppress replicative senescence of a *cdc13-F237A, E252K* mutant that disrupts both the Cdc13_EBM-N_ and Cdc13_EBM-C_ motifs (Fig 2D).

In summary, these findings indicate that mutation of *PIF1* allows sufficient telomerase recruitment to avoid replicative senescence caused by disruption of the Cdc13_EBM-C_-Est1 interaction alone, or double disruption of both the Cdc13_EBM-N_-Est1 and yKu-TLC1 interactions. However, suppression is not possible when both the Cdc13_EBM-C_-Est1 and yKu-TLC1 interactions are abolished. Disruption of both the Cdc13_EBM-N_-Est1 and Cdc13_EBM-C_-Est1 interactions can be suppressed by mutation of *PIF1* (Fig 2D), suggesting that the Cdc13_EBM-N_-Est1 interaction plays a more minor role, likely in support of the Cdc13_EBM-C_-Est1 interaction. Our findings suggest that Pif1 inhibits telomerase regardless of how telomerase is recruited: mutation of *PIF1* in *cdc13-2* cells allows increased telomerase recruitment via the yKu-TLC1 pathway, while mutation of *PIF1* in *cdc13-F237A tlc1Δ48* cells allows increased telomerase recruitment via the Cdc13_EBM-C_-Est1 pathway.

### Mutation of *PIF1* cannot suppress the replicative senescence of *est1Δ*

The *cdc13-2* mutation greatly reduces the recruitment of Est1 to telomeres (Chan *et al*., 2008), and expression of Cdc13-Est2 fusion protein allows cells to stably maintain their telomeres in the absence of Est1 (Evans & Lundblad, 1999). Therefore, it was possible that *pif1-m2* suppresses the replicative senescence caused by *cdc13-2* by somehow bypassing the need for Est1 for telomerase-mediated telomere extension. To test this idea, we sporulated an *EST1/est1Δ PIF1/pif1-m2* heterozygous diploid and monitored growth of the haploid meiotic progeny (Fig 2E). We find that *est1Δ pif1-m2* double mutants senesce like *est1Δ* single mutants, indicating that mutation of *PIF1* cannot bypass the need for Est1.

### Tel1 acts through the Cdc13_EBM-C_ motif to regulate telomere length

Since the *cdc13-2* mutation normally results in a complete defect in telomerase-mediated telomere extension, it has not been possible to perform classical genetic epistasis experiments to determine which telomere length regulators act through the Cdc13_EBM-C_-Est1 pathway. The viability and non-senescence of *cdc13-2 pif1-m2* strains gives us the opportunity to do so. The Rap1-interacting factors, Rif1 and Rif2, negatively regulate telomerase (Hardy *et al*, 1992; Wotton & Shore, 1997) while the Tel1 kinase is a positive regulator (Greenwell *et al*, 1995). We measured telomere length of haploid strains generated from the sporulation of heterozygous diploids (Fig 3). We find that *cdc13-2 pif1-m2* cells have short telomeres, which is in contrast to the more wild-type, but heterogeneous, length telomeres shown in Figure 1. The difference is most likely due to *cdc13-2* being expressed from a high-copy plasmid in Figure 1. While deletion of *RIF1* elongates *cdc13-2 pif1-m2* telomeres, both *cdc13-2 pif1-m2 rif2Δ* and *cdc13-2 pif1-m2 tel1Δ* triple mutants have very similar telomere lengths compared to *cdc13-2 pif1-m2*, indicating that Rif2 and Tel1 function upstream and in the same pathway as the Cdc13_EBM-C_ motif (Fig 3).

**Figure 3.**
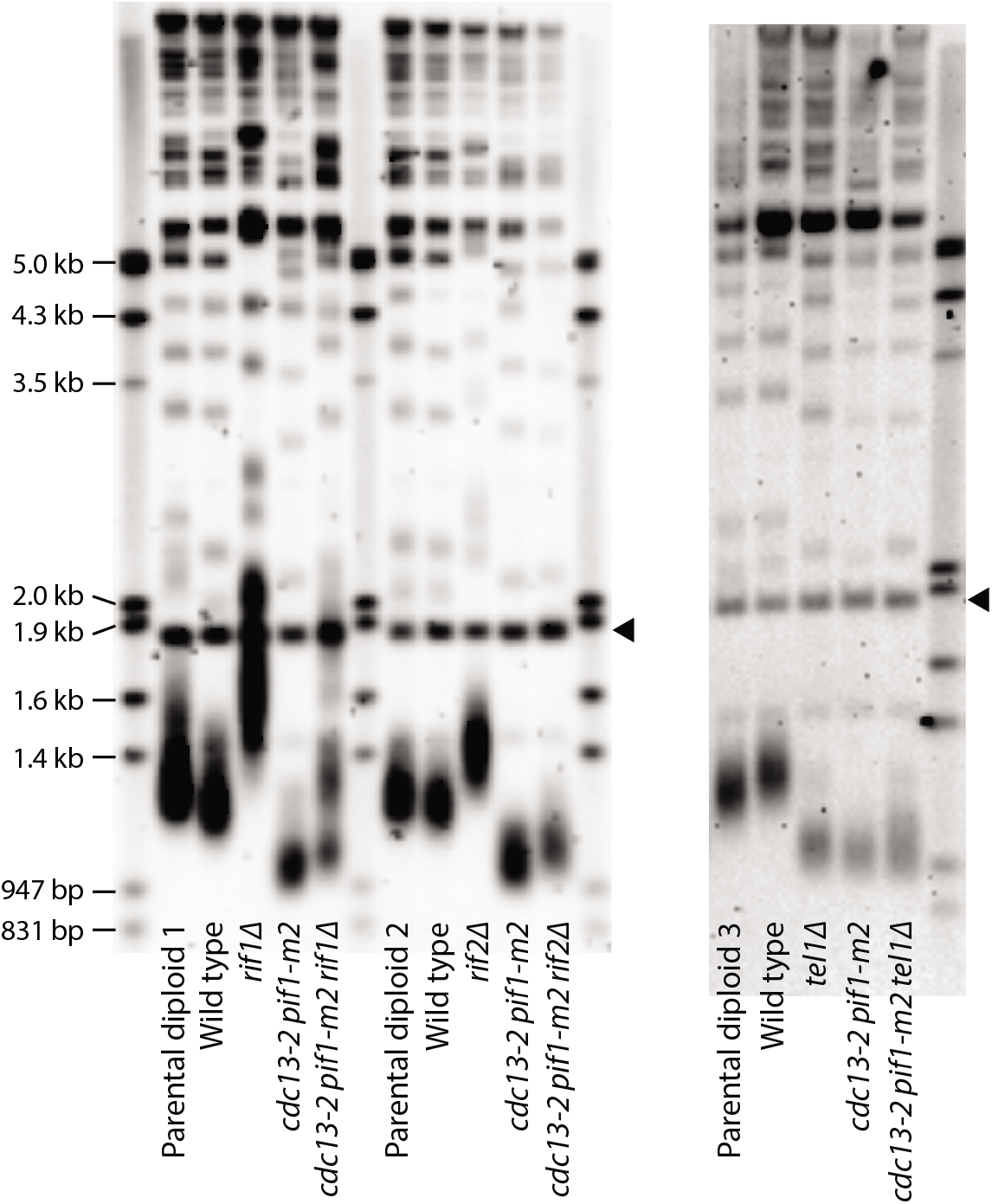
The *cdc13-2* allele is epistatic to *rif2Δ* and *tel1Δ* with respect to telomere length regulation in a *pif1-m2* background. Telomere Southern blot analysis of strains of the indicated genotypes. Parental diploids 1, 2, and 3 are EFSY8, EFSY9, and EFSY31, respectively. The black arrowhead indicates a 1.8 kb DNA fragment generated from the BsmAI-digestion of plasmid pYt103.

Tel1 often functions in concert with a related kinase, Mec1. Mutation of both *MEC1* and *TEL1* results in an *est* phenotype (Ritchie *et al*, 1999). Since Tel1 promotes telomerase activity through the Cdc13_EBM-C_-Est1 interaction, we examined whether the same is true for Mec1. If so, the replicative senescence of *mec1 tel1* double mutants should be suppressed by *pif1-m2*. We sporulated a *MEC1/mec1-21 TEL1/tel1Δ PIF1/pif1-m2* diploid strain and monitored the growth of the *mec1-21 tel1Δ* and *mec1-21 tel1Δ pif1-m2* haploid meiotic progeny. Both strains exhibited a similar rate of senescence (Fig 2F), indicating that Mec1 functions in a different pathway than Tel1 to promote telomerase activity, as previously proposed (Ritchie *et al*., 1999; Keener *et al*, 2019).

The nature of this Mec1-dependent pathway remains to be elucidated. Interestingly, association of Est1 and Est2 to telomeres is severely reduced by the deletion of *TEL1*, but largely unaffected by deletion of *MEC1* (Goudsouzian *et al*, 2006). Thus, it is unclear why *mec1 tel1* double mutants senesce. One possibility is that Mec1 and Tel1 redundantly phosphorylate a protein (or multiple proteins) to activate telomerase after it is recruited. There is evidence for telomerase activation after recruitment. For example, while expression of a Cdc13-Est2 fusion protein results in greatly elongated telomeres by forcing constitutive recruitment of telomerase to telomeres, hyperelongation is not seen in an *est1Δ* background, indicating that Est1 is important for both recruitment and activation of telomerase (Evans & Lundblad, 1999). Similarly, in addition to promoting the recruitment of telomerase, Tel1 affects telomerase activity by promoting the processive addition of telomeric repeats at critically short telomeres (Chang *et al*, 2007). Further work is needed to elucidate the exact mechanism by which Mec1 and Tel1 promote telomerase-mediated telomere extension. This is especially interesting considering that the mammalian homologs of Mec1 and Tel1—ATR and ATM, respectively—may also function in a similar manner (Lee *et al*, 2015).

## Material and Methods

### Yeast strains and plasmids

All yeast strains used in this study are listed in Table 1. Standard yeast genetic and molecular methods were used (Sherman, 2002; Amberg *et al*, 2005). Plasmids pEFS4 (pRS415-*cdc13-F237A*) and pFR96 (pRS415-*cdc13-F237A, E252K*) were created by site-directed mutagenesis of pDD4317 (pRS415-*CDC13*; Strecker *et al*., 2017) using primers designed by NEBaseChanger and the Q5 Site-Directed Mutagenesis Kit (New England Biolabs. Cat. No.: E0554S). The mutations were confirmed by DNA sequencing.

**Table 1.** Yeast strains used in this study.

| Strain name | Genotype | Source |
| --- | --- | --- |
| DDY3768 | *MATa·inc ura3·52 lys2·801 ade2·101 trp1·Δ63 his3·Δ200 leu2::natMX*<br>*rad52::HIS3 VII·L::TG34·HOcs·LYS2 ura3::hphMX cdc13::kanMX YEp24·*<br>*CDC13 pRS425·CDC13* | (Strecker *et al.*, 2017) |
| DDY3778 | *MATa·inc ura3·52 lys2·801 ade2·101 trp1·Δ63 his3·Δ200 leu2::natMX*<br>*rad52::HIS3 VII·L::TG34·Hocs·LYS2 ura3::hphMX cdc13::kanMX YEp24·*<br>*CDC13 pRS425·cdc13·E252K* | (Strecker *et al.*, 2017) |
| DDY3783 | *MATa·inc ura3·52 lys2·801 ade2·101 trp1·Δ63 his3·Δ200 leu2::natMX*<br>*rad52::HIS3 VII·L::TG34·Hocs·LYS2 ura3::hphMX cdc13::kanMX pif1·m2*<br>*YEp24·CDC13 pRS425·CDC13* | (Strecker *et al.*, 2017) |
| DDY3793 | *MATa·inc ura3·52 lys2·801 ade2·101 trp1·Δ63 his3·Δ200 leu2::natMX*<br>*rad52::HIS3 VII·L::TG34·Hocs·LYS2 ura3::hphMX cdc13::kanMX pif1·m2*<br>*YEp24·CDC13 pRS425·cdc13·E252K* | (Strecker *et al.*, 2017) |
| VSY20 | *MATa/MATα ade2·1/ADE2 can1·100/can1·100 his3·11, 15/his3·11, 15 leu2·*<br>*3, 112/leu2·3, 112 trp1·1/trp1·1 ura3·1/ura3·1 RAD5/RAD5 cdc13·*<br>*2::natMX/CDC13 pif1·m2/PIF1 yku80·135i::kanMX/YKU80* | This study |
| VSY7 | *MATa/MATα ade2·1/ADE2 can1·100/can1·100 his3·11, 15/his3·11, 15 leu2·*<br>*3, 112/leu2·3, 112 trp1·1/trp1·1 ura3·1/ura3·1 RAD5/RAD5 cdc13·*<br>*2::natMX/CDC13 pif1·m2/PIF1 tlc1Δ48::kanMX/TLC1* | This study |
| EFSY73 | *MATa/MATα ADE2/ADE2 can1·100/can1·100 his3·11, 15/his3·11, 15 leu2·*<br>*3, 112/leu2·3, 112 trp1·1/trp1·1 ura3·1/ura3·1 RAD5/RAD5*<br>*cdc13::kanMX/CDC13 pif1·m2/PIF1 tlc1Δ48::hphMX/TLC1 pRS415·cdc13·*<br>*F237A* | This study |
| FRY867 | *MATa/MATα ade2·1/ade2·1 can1·100/can1·100 his3·11, 15/his3·11, 15 leu2·*<br>*3, 112/leu2·3, 112 trp1·1/trp1·1 ura3·1/ura3·1 RAD5/RAD5*<br>*cdc13::kanMX/CDC13 pif1·m2/PIF1 tlc1Δ48::hphMX/TLC1 pRS415·cdc13·*<br>*F237A/E252K* | This study |
| CAY2 | *MATa/MATα ade2·1/ADE2 can1·100/can1·100 his3·11, 15/his3·11, 15 leu2·* | This study |
|  | *3, 112/leu2-3, 112 trp1-1/trp1-1 ura3-1/ura3-1 RAD5/RAD5*<br>*est1::HIS3/EST1 pif1-m2/PIF1* |  |
| MCY815 | *MATa/MATα ADE2/ADE2 can1-100/can1-100 his3-11, 15/his3-11, 15 leu2-3, 112/leu2-3, 112 trp1-1/trp1-1 ura3-1/ura3-1 RAD5/RAD5 mec1-21/MEC1 tel1ΔURA3/TEL1 pif1-m2/PIF1* | This study |
| EFSY8 | *MATa/MATα ade2-1/ADE2 can1-100/can1-100 his3-11, 15/his3-11, 15 leu2-3, 112/leu2-3, 112 trp1-1/trp1-1 ura3-1/ura3-1 RAD5/RAD5 cdc13-2::natMX/CDC13 pif1-m2/PIF1 rif1ΔHIS3MX/RIF1* | This study |
| EFSY9 | *MATa/MATα ade2-1/ADE2 can1-100/can1-100 his3-11, 15/his3-11, 15 leu2-3, 112/leu2-3, 112 trp1-1/trp1-1 ura3-1/ura3-1 RAD5/RAD5 cdc13-2::natMX/CDC13 pif1-m2/PIF1 rif2ΔHIS3MX/RIF2* | This study |
| EFSY31 | *MATa/MATα ade2-1/ADE2 can1-100/can1-100 his3-11, 15/his3-11, 15 leu2-3, 112/leu2-3, 112 trp1-1/trp1-1 ura3-1/ura3-1 RAD5/RAD5 cdc13-2::natMX/CDC13 pif1-m2/PIF1 tel1ΔURA3/TEL1* | This study |

### Liquid culture senescence assay

Liquid culture senescence assays were performed essentially as previously described (van Mourik *et al*, 2016).

### Telomere Southern blot

Telomere length analysis by Southern blotting was performed essentially as previously described (van Mourik *et al*, 2018). A 1.8 kb DNA fragment containing telomeric sequences generated from the BsmAI-digestion of plasmid pYt103 (Shampay *et al*, 1984) was loaded together with each sample. Southern blots were probed with a telomere-specific probe (5’-TGTGGGTGTGGTGTGTGGGTGTGGTG-3’).

## Acknowledgments

We thank V. So for help with strain construction, and A. Bertuch for providing the *tlc1Δ48* and *yku80-135i* mutants.

## Author contributions

Conceptualization: EFS, MC; Investigation: EFS, FRRB, SS, MC; Writing – original draft: EFS, MC; Writing – review & editing: EFS, FRRB, SS, MC; Supervision: MC.

